# Longer Interstimulus Intervals Enhance Efficacy of Automated Phase-Targeted Auditory Stimulation on Procedural Memory Consolidation

**DOI:** 10.1101/2024.12.26.630252

**Authors:** Vanessa Kasties, Nicole Meier, Nora-Hjördis Moser, Renske Sassenburg, Walter Karlen, Maria Laura Ferster, Sara Fattinger, Angelina Maric, Reto Huber

## Abstract

Up-phase-targeted auditory stimulation (up-PTAS) during slow-wave sleep has become a valuable tool for modulating slow oscillations and slow-oscillation-spindle-coupling in favor of overnight memory retention. Developing effective, automated protocols for translation into more naturalistic or clinical settings is an ongoing challenge, especially given that current PTAS protocols and their behavioral effects vary greatly between different studies. Here, we assessed the electrophysiological and behavioral effects of systematically varying interstimulus intervals (ISIs) in automated up-PTAS in the home setting, using a mobile PTAS device and app-based behavioral tasks. Building on studies suggesting a non-linear relationship between stimulus number and PTAS effects, we show that applying fewer stimuli with longer ISIs enhanced overnight memory consolidation of a finger-tapping sequence more effectively than applying more stimuli with shorter ISIs. The behavioral response was predicted by the number of stimuli with auditory evoked K-complexes relative to the number of stimuli without K-complexes. PTAS stimuli applied at longer ISIs (> 1.25) were associated with a higher likelihood of K-complex responses and fast spindles nesting in the K-complex up-phase. Our results suggest that up-PTAS can be optimized for overnight memory consolidation by introducing ISIs of at least 1.25s. Our study highlights the feasibility of longitudinal at-home PTAS combined with app-based behavioral tasks in healthy participants while leveraging the mechanistic insights such data can offer.

**Statement of Significance:** Phase-targeted auditory stimulation (PTAS) holds great promise for non-invasively enhancing essential functions of slow-wave sleep. However, current protocols have produced variable results and are often confined to laboratory settings. Our study demonstrates the feasibility of a mobile application of automated PTAS and provides experimental evidence that prolonging interstimulus intervals positively affects overnight procedural memory consolidation via K-complexes and coupled sleep spindles. As K-complexes may also be involved in cardiovascular function and brain waste clearance, the proposed protocol optimization may have effects beyond memory enhancement. Together, our findings lay a foundation for a broader application of PTAS in clinical longitudinal studies to improve patient care and recovery outcomes.

## INTRODUCTION

Sleep is essential to our health and well-being. Therefore, it is no surprise that sleep monitoring and modulation techniques are gaining popularity. As mobile recording devices have become more reliable and technically sophisticated^1–5^, high-quality sleep data can be obtained in the home setting, offering high ecological validity, cost-effectiveness, and user comfort. Home-based sleep studies reduce the burden for participants, as they are not required to travel or change their normal sleeping environment. In combination with wearable recording devices that the participants can apply independently, the home setting provides ideal conditions for sleep monitoring over extended periods^2,6,7^. Thus, mobile technology holds great promise for clinical longitudinal research^8^. However, ensuring its accessibility and physiological validity in healthy controls is an important next step to advancing its future clinical applicability.

Up-phase targeted auditory stimulation (up-PTAS) has become a valuable tool for modulating sleep non-invasively without disrupting the natural sleep architecture^9,10^. Up-PTAS relies on applying short (50 ms) bursts of pink noise at the up-phase of ongoing slow oscillations^11^. This has been shown to boost the sleep slow (< 1 Hz) oscillation, which orchestrates thalamic spindles (oscillatory bursts of 11 – 16 Hz) and hippocampal ripples to the depolarizing up-phase^11^. This nesting of sleep oscillations is assumed to be critical for memory consolidation^12,13^. Indeed, the enhancement of low (0.5 – 1 Hz) slow wave activity (SWA), along with changes in spindle power, was directly associated with improved overnight declarative memory consolidation in studies applying up-PTAS^11,14^. Thus, in principle, PTAS holds great promise for boosting the memory function of sleep, from which clinical populations may profit. However, there seem to be inconsistencies between studies. A recent meta-analysis has highlighted that up-PTAS positively affects declarative memory overall, but the effect sizes are small and vary considerably across studies^15^. Moreover, the few studies that examined other domains, such as procedural memory or executive functions, offer conflicting evidence^16–19^. These inconsistencies may be caused by variations in PTAS protocols leading to different evoked electrophysiological and behavioral responses. This underlines the necessity for systematic comparisons of PTAS protocols to determine the impact of different stimulation parameters.

The two studies comparing up-PTAS protocols systematically found no advantage of continuous stimulation over intermittent (e.g., stimulation in isolated pairs with fixed interstimulus intervals, stimulation in ON-OFF windows) approaches on low SWA enhancement or memory retention, suggesting a self-limiting mechanism^6,14^. Along these lines, a recent study showed that up-phase targeted stimuli played in short succession rather induce a long SWA plateau, whereas stimuli preceded by 6 s OFF windows reliably evoke a K-complex response, irrespective of the targeted phase^20^. This observation matches early reports of evoked K-complexes being susceptible to habituation or refractory processes^21–23^. Interestingly, the K-complex response was accompanied by evoked spindles that were preferentially nested in the up-phase of the 1 Hz oscillation and were associated with improved declarative memory performance^20^. Coagulating these findings, PTAS for memory consolidation may benefit from an approach that considers K-complex refractory periods, which should allow fewer stimuli to evoke full electrophysiological responses. Therefore, we propose interstimulus interval (ISI) as an important stimulation parameter in PTAS, the effects of which were evaluated in this study.

We conducted a study in the home setting, using a mobile electroencephalography (EEG) device, the Mobile Health Systems Lab *SleepBand* v3 (MHSL-SB, ETH Zurich, Zurich, Switzerland), to apply two up-PTAS protocols with long or short ISIs and testing their effects on the consolidation of a tablet-based finger tapping task. The finger-tapping task (FTT) is an explicit motor task that has collectively been reported to profit from sleep-dependent consolidation^24^. It was particularly suited for the home setting because of its high level of standardization and automatization. We hypothesized that using this setup, our experimental at-home up-PTAS protocol combined with an app-based FTT would be both feasible and valid, replicating previous in-lab findings. Furthermore, supported by previous literature^6,14^, we hypothesized that the long ISI up-PTAS protocol would drive behavioral responses more efficiently (i.e., producing at least similar effects with fewer stimuli) than the short ISI protocol. To gain a mechanistic understanding, we explored how these systematic variations in the ISI influenced electrophysiological effects, focusing specifically on stimulus-evoked K-complexes. Our results confirm the feasibility of PTAS studies in the home setting and provide exploratory mechanistic insights into the benefits of prolonging the interstimulus interval for enhancing procedural memory consolidation.

## METHODS

### Participants

Eighteen healthy young adults (mean age: 27 ± 5.16 years, eight male) were recruited through an online advertisement on a university platform and word of mouth from March to December 2022. Eligible candidates answered an anonymous questionnaire that tested them for the following inclusion criteria: German proficiency of at least C1 level, right-handedness, good self-reported sleep and a regular sleep-wake cycle, good general health, and ability to comprehend and adhere to study procedures. Candidates were excluded if they reported psychiatric, neurologic, or physical disorders, hearing or uncorrected vision impairments, skin allergies, shift work, or had traveled through more than one time zone in the past month. Furthermore, candidates were excluded if they engaged in gaming more than four times a week, were professional musicians, or had previously participated in similar sleep studies. These measures were taken to reduce practice effects on the study tasks. Participants selected for the study agreed to limit caffeine consumption to maximally two standard portions per day, abstain from alcohol during assessment days, and from cannabis or other drugs throughout the study. All participants gave written informed consent. The study protocol followed the guidelines provided by the Declaration of Helsinki and was approved by the cantonal ethics committee (BASEC2021-02486).

### Experimental protocol

The protocol comprised an adaptation phase and two intervention nights separated by one week, as shown in **Figure 1**. All study procedures were performed in the home setting. During the adaptation phase, a study member visited the participants at home to train them in the study tasks and procedures. Following the home visit, participants underwent one to two adaptation nights with a mobile, wearable PTAS device (MHSL-SB). The adaptation phase was followed by a wash-out phase of approximately one week. On intervention nights, two stimulation protocols were applied in random order. The two intervention nights had identical timelines. In the evening, starting two hours before bedtime, participants completed a diary, the Karolinska Sleepiness Scale^25^ (KSS), the psychomotor vigilance task^26^ (PVT), and the FTT, presented within a tablet-based test battery. A digital checklist helped them mount the PTAS device. In the morning, participants removed the device aided by a digital checklist, completed a sleep diary and the KSS once again, and performed retrieval tests for the FTT and word-pair tasks. To reduce the impact of sleep inertia, participants were asked to wait at least 30 minutes after awakening before completing their tasks. All nights were scheduled according to the participants’ habitual sleep and wake times. Participants’ adherence to the protocol was ensured by detailed schedules and information material, regular phone calls, and remote data monitoring.

**Figure 1:**
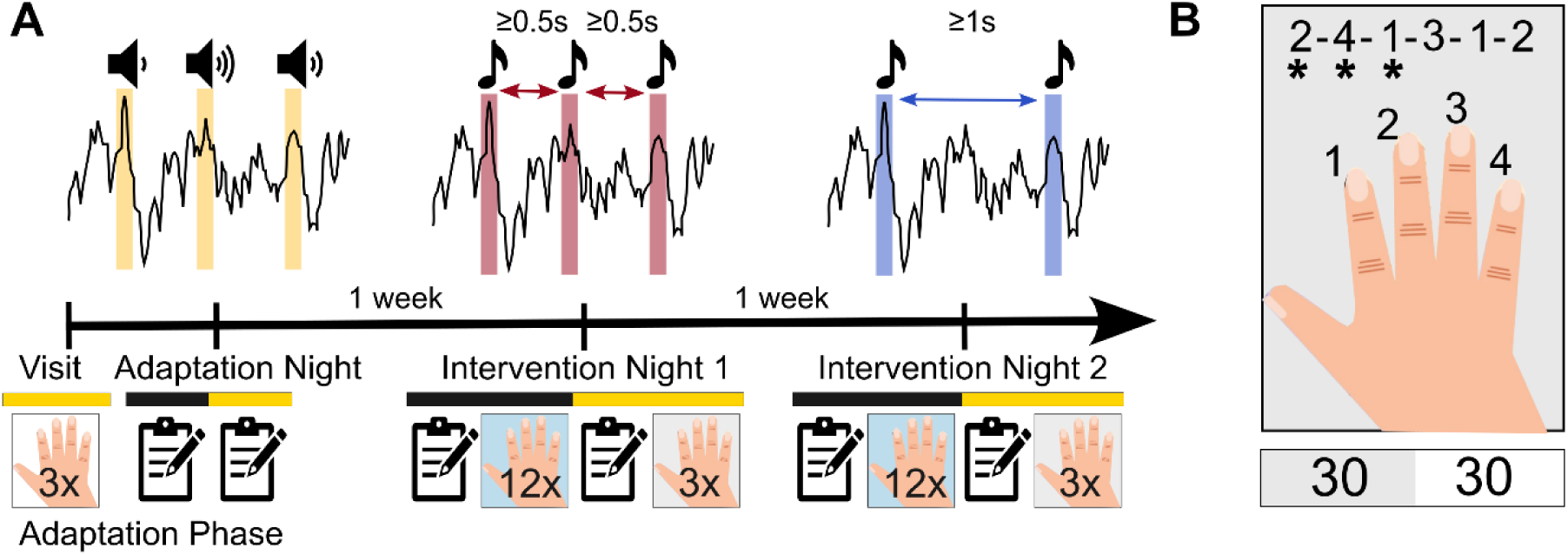
Study and task protocols. **(a)** One representative participant’s timeline. Icons represent the finger tapping task with the respective number of blocks (backgrounds: white - training, blue - learning, grey - retrieval) and questionnaires surrounding the stimulation nights. Black bars mark tasks completed in the evening, and yellow bars mark tasks completed in the morning. During the Adaptation Night, the optimal stimulation volume was determined. During Intervention Night 1 and Intervention Night 2, a long ISI (ISI ≥ 1 s) or a short ISI (ISI ≥ 0.5 s) up-phase targeted auditory stimulation (up-PTAS) protocol was administered in consecutive windows of 6 s ON and 6 s OFF. **(b)** Schematic representation of tablet-based finger tapping task, where asterisks tracked the current position in the sequence in real-time. A representative task block is shown below, where numbers denote seconds and colors active (grey) or break (white) periods.

### Behavioral assessments

Procedural memory consolidation was assessed using a tablet version of the FTT, adapted from Karni et al., 1998^27^. In this task, participants were shown a six-digit sequence on the screen that they were instructed to tap with their dominant hand as fast and accurately as possible. The numbers (1 – 4) in the sequence corresponded to the index, middle, ring, and pinky finger. The sequence was displayed in the upper half of the screen throughout the task, and a marker indicated where in the sequence participants were. Sequences were never used twice in the same participant, randomized across stimulation conditions, and matched in difficulty. Participants trained in an easy sequence (1 – 2 – 3 – 3 – 4 – 4) for three trials during the adaptation visit to familiarize themselves with the task. Each trial comprised blocks of 30 s tapping and 30 s break to prevent motor fatigue. In the evening session, participants practiced a novel sequence (2 – 4 – 1 – 3 – 1 – 2 or 2 – 1 – 3 – 1 – 4 – 2) for twelve trials, using the same block-wise design. The same sequence was tested in three retrieval trials in the morning session. We quantified performance and variability measures like previous reports^28–31^. Performance per trial was calculated as the percentage of correct sequences divided by the intertap interval in seconds. This marker accounts for the speed-accuracy tradeoff during skill learning^32,33^. Tapping variability was determined as the average standard deviation of intertap intervals in completed sequences per trial^28,31^.

To assess objective and self-reported vigilance, participants completed a standard ten-minute PVT and the KSS in the evening and the morning around stimulation nights. In the PVT, participants were presented with a red rectangular box on the tablet screen and were instructed to tap on the screen as soon as a yellow stimulus counter appeared. When a response was recorded, the counter stopped to display the reaction time (RT) in milliseconds. We assessed the response speed (1/RT) in non-lapse trials (RT < 500 ms) as a marker of sustained attention.

### Sleep recordings

EEG, electrooculography (EOG), and electromyography (EMG) were recorded with the MHSL-SB. The frontal EEG channel (Fpz), two EOG channels, and two EMG channels were referenced against the right mastoid, whereas the left mastoid served as a ground electrode. Electrodes were single-use and auto-adhesive (Neuroline 720, Ambu A/S, DK). Their impedance was assessed automatically at the beginning of every recording and checked post-hoc during data quality assessments (average impedance: 90.17 ± 69.45 kOhm). Signals were recorded at a 250 Hz sampling rate using a 24-bit analog-to-digital converter, subjected to a real-time anti-aliasing filter and a 50 Hz notch filter, and piped to the built-in stimulation algorithm. The algorithm categorizes the EEG into NREM and not-NREM sleep based on spectral power thresholds in the low delta (2 – 4 Hz), high delta (3 – 5 Hz), and high beta (20 – 30 Hz) bands^4^. We derived a set of default power thresholds based on historical data of a similar age cohort, which we used throughout the stimulation nights. Upon detection of stable NREM sleep for ten minutes, stimulation is enabled. A phase-locked-loop algorithm^4^ is then employed to detect slow waves in real-time and trigger stimuli at the target phase of 45°, corresponding to the down-to-up transition of the ongoing slow wave. We configured the device to stop the stimulation 2.5 hours after the first stimulus to leave the second half of the night untouched. This procedure was introduced to mitigate effects of PTAS on REM sleep, which has been reported to affect mood^6^. Stimulation was applied in a block design of 6 s ON windows, where stimulation was enabled, and 6 s OFF windows, where stimulation was disabled. Stimulation flags in OFF windows (sham stimuli) were reconstructed offline based on the algorithm’s power thresholds. The stimuli were 50 ms bursts of pink noise applied via headphones integrated into a headband.

### Adaptation night stimulation protocol

Given previous reports that found a correlation between stimulus response and memory consolidation after PTAS^34^, we leveraged the adaptation nights to individualize the stimulation volume to a level that achieved quantifiable PTAS responses without arousing the participant. If the optimal stimulation volume could not be determined after one adaptation night, another night was recorded. The volume determined in the adaptation phase was used constantly throughout the intervention nights. The final volumes ranged between 46 dB sound pressure level and 64 dB sound pressure level.

### Intervention night stimulation protocols

During intervention nights, two up-PTAS protocols were administered in random order: the short ISI protocol and the long ISI protocol. Both protocols shared the same stimulation period, an ON-OFF block design, and a constant stimulation volume as derived from the adaptation phase but differed systematically in the interstimulus intervals. In the short ISI protocol, we selected an interstimulus interval of ≥ 0.5 s, with the expectation of stimulating every detected slow wave within ON windows once. In the long ISI protocol, we extended the interstimulus interval to ≥ 1 s, with the expectation of skipping certain slow waves within a train or only stimulating consecutive waves in the slow oscillation frequency range (≤ 1 Hz).

### Sleep scoring and artifact rejection

For sleep scoring, signals were preprocessed and scored in *Visbrain*^35^ (running in Python 3.8.10) according to AASM guidelines^36^. To filter the signal, 3rd-order Butterworth digital IIR filters were applied forward and backward to avoid phase shifts. The signal from Fpz was band-pass filtered between 0.5 Hz and 35 Hz. The signals from the two eye channels were band-pass filtered between 0.3 Hz and 35 Hz. Signals from the two muscle channels were band-pass filtered between 10 Hz and 100 Hz and re-referenced bilaterally if visually assessed signal quality was good in both channels. After filtering, all signals were downsampled to 128 Hz, as required by the scoring software. Vigilance states (wake, N1, N2, N3, REM) were visually scored by two sleep experts in 30 s epochs in *Visbrain*^35^ and validated by a third sleep expert. All scorers were blind to the experimental condition and the stimulation flags. After scoring, NREM (N1, N2, N3) sleep epochs were subjected to a semi-automatic artifact detection routine adapted from Leach et al., 2020^37^ for single-channel signals.

### Electroencephalography signal preprocessing

The EEG signal was first subjected to a low-pass filter with a pass band of up to 30 Hz (−6 dB attenuation at 39.86 Hz; filter order: 46, at a sampling rate of 250 Hz), then subjected to a high-pass filter with a pass band starting at 0.5 Hz (−6 dB attenuation at 0.37 Hz; filter order: 2988, at a sampling rate of 250 Hz), similar to a previous publication from our laboratory^20^. Filters were Kaiser window FIR filters with zero-phase shift to avoid phase distortions. Filtering and all consecutively described signal processing steps were performed in MATLAB R2022a (The MathWorks Inc., 2022) using EEGLAB^38^ and the FieldTrip toolbox^39^.

### Analysis of slow-wave activity response

Average SWA in the low (0.75 – 1.25 Hz) and adjacent (1.25 – 4 Hz) frequency bands were calculated separately in ON and OFF windows, and the absolute difference between ON and OFF was obtained as a marker of stimulation response^6,40,41^. To this end, we selected only ON windows in scored NREM sleep (stages N2 and N3) that contained at least one stimulus and their consecutive OFF windows. SWA was calculated with MATLAB’s *pwelch* function for each 6 s window, using 4 s Hanning windows with 50 % overlap, and then averaged separately across all ON and OFF windows.

### Quantification of K-complexes

Stimulus-associated K-complexes were detected in the preprocessed Fpz signal using the *DETOKS* algorithm^42^. Briefly, the algorithm first removes transients from the signal and then splits it into a low-frequency component and an oscillatory component. A Teager-Kaiser energy operator (TKEO) is applied to the low-frequency component for detecting K-complexes and to the oscillatory component for detecting spindles. TKEO thresholds must be manually adjusted and were optimized for this data set to obtain the maximal number of events in NREM epochs but a minimum number of events in artifactual epochs.

Before subjecting the Fpz signal to the *DETOKS* algorithm for K-complex detection, we automatically removed pulse artifacts from the data using a published algorithm for single-channel data (*brMEGA*)^43^. The low-pass filter cutoff for the low-frequency component was left at the default value (4 Hz), but we extended the duration threshold for K-complexes to 0.5 – 2 s (corresponding to a frequency range of 0.5 – 2 Hz) to be comparable to previous reports^44,45^ Stimulus-associated K-complexes were defined to start within 1.5 s after stimulus onset and were determined separately for ON stimuli and sham stimuli. The final detection result was visually validated in individual trials by plotting the preprocessed signal against the binary detection time series. Furthermore, we assessed the average waveform and time-frequency response across detected events (2 s epochs, aligned at K-complex start) in comparison to a random selection of non-labeled NREM sleep trials. Time-frequency responses were assessed using Morlet wavelets ranging from 1 – 30 Hz using the procedure described in *Time-frequency analysis*.

### Determination of ISI bins

To determine in more detail at which interstimulus interval stimulus-associated K-complexes were most likely to occur, we defined ISI bins using a data-driven approach. Defining such a uniform scale was necessary because ISIs were not fixed across the sample but varied highly depending on the underlying slow waves. ISIs preceding auditory stimuli were determined across the sample. Splitting the distribution into nine quantiles and rounding the quantile edges down to the nearest 0.5 s defined an initial set of seven bin edges. After removing duplicates, these edges were further subdivided into smaller bins between 0.5 s and 2.5 s, as this was the range where we expected the most change to happen. This procedure produced a final set of ten ISI bins: 0.5 – 0.75 s, 0.75 – 1 s, 1 – 1.25 s, 1.25 – 1.5 s, 1.5 – 2 s, 2 – 2.5 s, 2.5 – 6.5 s, 6.5 – 8 s, 8 – 10.5 s, >10.5 s. We then collapsed all available nights per individual and quantified the number of stimuli followed by a K-complex and the number of stimuli followed by no K-complex per ISI bin for each participant, defining event vs. no event as a binary outcome.

### Time-frequency analysis

We set the ISI bin where K-complex probability plateaued as the threshold to categorize stimuli based on their preceding ISI. The data was then segmented into 4 s epochs around each stimulus, with a pre-stimulus time of 1s, using FieldTrip’s *redefinetrial* function. These epochs surrounding ON stimuli (ON epochs) were contrasted with epochs surrounding matching sham/ OFF stimuli (OFF epochs).

The auditory evoked potential (AEP) of each stimulus category was determined by averaging across all ON and OFF epochs separately and then calculating the difference between ON and OFF. The time-frequency response of each stimulus category was obtained by convolving the signal in ON and OFF epochs with a range of Morlet wavelets covering a linearly increasing frequency space from 1 Hz to 30 Hz in 1 Hz steps. Cycle number increased linearly as a function of frequency and ranged from 3 to 17.5 cycles, with a step width of 0.5 cycles per frequency bin. Average time-frequency power maps were log-normalized by a whole-epoch baseline calculated separately for each frequency bin across all ON and OFF epochs. After that, the difference map (ON-OFF) was computed for visualization.

To assess statistical differences between ON and OFF epochs, we used a nonparametric clustering procedure^46^ with the following specifications: First, paired Student’s t-tests were computed (two-tailed, α = 0.05) for the difference between ON and OFF for all pixels of the time-frequency map, resulting in a real t-map. Then, reference t-maps were obtained by randomly permuting ON and OFF epochs within each night and performing a paired t-test on the permuted time-frequency maps. The permutation was repeated 2000 times to obtain a reference distribution for each pixel. Cluster thresholds were defined based on the 95th percentile of clusters of significant t-values (two-tailed, α = 0.05) in reference maps. Lastly, this cluster threshold was applied to the real t-map to obtain significant clusters. An equivalent cluster correction was applied to the t-time-series resulting from ON-OFF AEP contrasts (paired two-tailed Student’s t-tests, α = 0.05). Cluster permutation statistics were performed in MATLAB.

For a hypothesis-driven assessment of the differences between the stimuli below and above the ISI threshold, we extracted features from the ON-OFF contrasts in time and time-frequency space. The prominent N550 AEP component was quantified for each stimulus category and night in two ways: firstly, by calculating the mean average in the time window from 500 to 600 ms and secondly, by quantifying the negative area under the curve within a broader time window of 200 - 1000 ms^47^. Time windows were visually determined based on the grand average ON-OFF AEP, collapsed across stimulus categories. To ensure more stable mean estimates of AEP variables, we only included contrasts where at least 50 epochs were obtained for ON and sham stimuli, respectively. From the grand average time-frequency map, we visually determined regions of interest in the delta, theta, alpha, and sigma bands. We extracted the average baseline normalized power from these regions for each stimulus category and night. These features were then used for comparisons between stimulus categories.

### Phase-amplitude modulation analysis

To assess the coupling strength between the fast spindle (13 – 16 Hz) band and the underlying wave in the slow oscillation frequency range (1 Hz) within 2 s after the stimulus, we computed the z-scored modulation index (zMI)^48^ separately per night and stimulus category. The z-scoring mitigates confounding by the absolute power of the power-providing signal, which may affect contrasts between different sets of epochs or conditions^49^. For the z-transformation, we constructed a null distribution of MI by repeatedly (n = 200) cutting each sigma-epoch at a random sample and reversing the order of both parts^50^. The empirical MI was then z-transformed with the null distribution, resulting in an MI measure that is independent of power differences between stimulus categories. For this analysis, we used 18 phase bins of 20°, similar to previous studies^48^. The frequency range for the sigma band was determined based on previous reports of fast spindles being preferentially coupled to the up-phase of slow oscillations and, therefore, potentially relevant for sleep-dependent memory processes^51^. The frequency of the underlying wave was determined to match the frequency ranges reported for K-complexes^52,53^ and previous reports that found most spindle coupling in the up-phase of the 1 Hz oscillation^20,54^.

### Statistics

Statistical analyses included paired parametric or nonparametric tests for comparing variables between the stimulation conditions or stimulus categories and robust linear mixed effects models for assessing the linear relationship between variables of interest and behavioral outcomes. For paired comparisons, we assessed the residuals by visually examining the histogram and Q-Q plot and formally testing their distribution for normality with a Shapiro-Wilk-Test. If the criteria for normal distribution were met, the comparison was performed with a two-sided Student’s t-test. Otherwise, we chose a nonparametric two-sided Wilcoxon signed-rank test. Linear mixed effects models always included random intercepts for participants. Model fit was assessed for homoscedasticity, linearity, and independence of residuals, normal distribution of residuals and random effects, and influential points. As influential points were present in most models, we chose to fit robust linear mixed models using the *robustlmm* package in R^55^. For correlations concerning only one condition, we used robust percentage bend correlations^56^. For all statistical tests, p-values below 0.05 were considered significant. In exploratory analyses, we report p-values below 0.1 as trends. P-values for robust linear mixed models were approximated based on the t-statistic and degrees of freedom reported in the model output. Unless stated differently, all statistical analyses were performed in R Statistical Software (v4.4.1; R Core Team, 2024).

## RESULTS

### Feasibility of automated auditory stimulation in the home setting

We conducted a randomized cross-over study, subjecting 18 healthy participants to two different up-PTAS protocols and a series of behavioral tasks in the home setting. Adherence to the protocol was monitored in several ways. Firstly, data quality was assessed daily after the data was automatically transmitted from the participant’s device to the server. Secondly, time stamps of task completion were compared to the participants’ individual schedules. Overall, 16 participants had complete EEG data from both intervention nights, and data from these participants were used for EEG analyses that did not involve associations with FTT data. Of this subsample, 14 participants had complete FTT data for both intervention nights. The EEG data from the long ISI stimulation night was lost in two participants for technical reasons. One participant was excluded from behavioral analyses of the FTT due to completing the learning and retrieval tasks in the wrong order, and one participant was excluded for deviating from their task schedule by several hours after one intervention night but not the other. In summary, most participants adhered well to the protocol, where EEG data loss could be solely attributed to technical reasons.

First, we assessed the impact of the short ISI and long ISI PTAS protocols on self-reported sleepiness on the KSS and standard sleep architecture variables, as well as next-day vigilance, assessed by PVT speed (1/RT). Of the 16 participants with complete EEG data, KSS data was obtained from 15 individuals, and complete PVT data from 13 individuals. The different stimulation protocols did not show any differential effects on sleep architecture, PVT, or KSS (**Table 1**), except for a significantly higher percentage of N1 sleep in the long ISI condition compared to the short ISI condition (*g = 0.48, p = 0.04*). In summary, we found no differential effects of the two protocols on self-reported variables, and objective sleep markers were largely similar.

**Table 1:** Comparison of sleep architecture, KSS and PVT outcomes between the stimulation conditions.

|  | *Short ISI*<br>*(mean ± SD)* | *Long ISI*<br>*(mean ± SD)* | *Mean*<br>*Difference* | *95% CI* | *g* | *p* |
| --- | --- | --- | --- | --- | --- | --- |
| *TTB [min]* | 468 ± 46.6 | 485 ± 6.7 | 17.3 | [-15.0; 49.5] | 0.39 | 0.27 |
| *TST [min]* | 399 ± 54.8 | 396 ± 46.4 | -3.4 | [-24.5; 17.7] | -0.06 | 0.73 |
| *SL [min]* | 31 ± 23.2 | 39 ± 46.0 | 7.9 | [-7.4; 23.3] | 0.14 | 0.29 |
| *SEF [%]* | 89.8 ± 6.38 | 87.9 ± 9.88 | -1.90 | [-4.62; 0.83] | -0.17 | 0.16 |
| *WASO [min]* | 14 ± 9.6 | 24 ± 30.4 | 9.6 | [-5.4; 24.6] | 0.36 | 0.19 |
| *REML [min]* | 79 ± 26.9 | 104 ± 57.0 | 25.0 | [-1.8; 51.8] | 0.49 | 0.07 |
| *N1 [%]* | 4.7 ± 2.05 | 6.0 ± 2.87 | 1.32 | [0.11; 2.53] | **0.48** | **0.04** |
| *N2 [%]* | 45.0 ± 7.82 | 42.0 ± 8.16 | -2.98 | [-6.79; 0.83] | -0.35 | 0.12 |
| *N3 [%]* | 19.7 ± 5.87 | 18.9 ± 4.79 | -0.77 | [-2.94; 1.40] | -0.13 | 0.46 |
| *REM [%]* | 20.5 ± 4.79 | 21.0 ± 6.34 | 0.54 | [-2.11; 3.19] | 0.09 | 0.67 |
| *Wake [%]* | 10.2 ± 6.39 | 12.1 ± 9.88 | 1.89 | [-0.83; 4.62] | 0.17 | 0.16 |
| *ΔKSS* | -0.5 ± 1.77 | -0.2 ± 2.18 | 0.27 | [-0.79; 1.32] | 0.13 | 0.60 |
| *Morning KSS* | 3.4 ± 1.30 | 3.8 ± 2.04 | 0.4 | [-0.53; 1.33] | 0.21 | 0.37 |
| *ΔPVT* | 0.13 ± 0.20 | 0.06 ± 0.11 | 0.070 | [-0.02; 0.16] | 0.38 | 0.13 |
| *Morning PVT* | 2.84 ± 0.36 | 2.79 ± 0.35 | 0.059 | [-0.01; 0.13] | 0.16 | 0.09 |

Mean differences refer to the contrast long ISI > short ISI. The reported p-values were obtained by paired Student's t-tests (N = 16) and effect size is reported in terms of Hedge's g. Significant comparisons are marked in bold. Abbreviations: TTB – total time in bed, TST – total sleep time, SL – sleep latency, SEF – sleep efficiency, WASO – wake after sleep onset, REML – rapid eye movement sleep latency, N1 – NREM sleep stage 1, N2 – NREM sleep stage 2, N3 – NREM sleep stage 3, REM – rapid eye movement sleep, ΔKSS – overnight change in Karolinska Sleepiness Scale, ΔPVT – overnight change in psychomotor vigilance task.

To evaluate whether the ISI manipulation was successful, we quantified the stimulation characteristics in each condition. The overall number of stimuli applied per night ranged from 186 to 2028, with an average of 1098 ± 501 in the short ISI condition and 638 ± 260 in the long ISI condition. The total number of stimuli was significantly lower in the long ISI condition compared to the short ISI condition (*g = −0.98, p < 0.001*). Accordingly, the long ISI condition showed a significantly longer median interstimulus interval within ON windows (long ISI: mean = 1.92 ± 0.13s, short ISI: mean = 1.15 ± 0.14s, long ISI > short ISI: *g = 5.36, p < 1*10^-9^*). Together, these data show that the two protocols were clearly distinguishable based on their interstimulus intervals.

### Effects of auditory stimulation differences on procedural task performance

Participants were assessed on the FTT before and after each stimulation night to determine the effect of the two auditory stimulation protocols on procedural memory consolidation. They completed twelve learning trials of a novel sequence in the evening and three retrieval trials of the same sequence the following morning. We quantified procedural learning with a performance score, which accounts for the speed-accuracy tradeoff, and tapping variability, as in previous reports^28–31^.

**Figure 2A** shows that participants exhibited a logarithmic learning trajectory during the training session. Their performance increased rapidly during the first three trials and then approached a plateau. Likewise, participants exhibited an inverse logarithmic learning curve for tapping variability, with rapid reductions during the first three trials and a plateau in later trials. The learning plateau for each participant and condition was approximated by the average of the last three learning trials (10 – 12) in the evening session. The learning rate was approximated by the difference between the first trial, considered the baseline, and the learning plateau. Neither performance nor tapping variability differed significantly between conditions at baseline (performance: *g = 0.24, p = 0.19;* tapping variability: *g = −0.06*, *p = 0.81*). Tapping variability learning rate and plateau were also not significantly different between conditions (plateau tapping variability: *g = 0.31, p = 0.18;* learning rate tapping variability: *g = 0.30, p = 0.33)*. Robust linear mixed models controlling for period and period×condition interaction also indicated no significant main effect of condition on any learning variable related to tapping variability (baseline: *β = −2.89, t = −0.11, p = 0.90;* plateau: *β = 17.98, t = 1.22, p = 0.23*; learning rate: *β = 26.38, t = 1.04, p = 0.31)*. However, we found significant differences in plateau performance and learning rate (plateau performance: *g = −0.41, p = 0.04*; learning rate performance: *g = −0.67*, *p = 0.03*). The main effect of condition remained significant when controlling for period and period×condition interaction in robust linear mixed models (baseline performance: *β = 27.08, t = 2.67, p = 0.01*, plateau performance: *β = −33.93, t = −2.31, p = 0.03*; learning rate performance: *β = −56.12, t = −3.72, p = 0.001*). These results indicate that at the stage of learning, tapping variability evolved similarly across conditions, whereas performance exhibited differences between conditions already before the PTAS intervention. To exclude the possibility of spurious results driven by differences at baseline, we decided to exclude performance-related measures from further analysis.

Next, we assessed whether overnight memory consolidation differed between the stimulation conditions. Overnight memory consolidation was defined as the difference between the learning plateau and the average of the three retrieval trials, as established in prior work^57^. The reduction in overnight tapping variability was significantly greater in the long ISI condition than in the short ISI condition (*g* = −0.64, *p* = 0.04; **Figure 2B**). This effect persisted when controlling for period and period×condition interaction (main effect of condition: *β = −32.36, t = −2.13, p = 0.04*). Notably, the long ISI stimulation benefited procedural memory consolidation despite the lower number of stimuli compared to the short ISI stimulation. To exclude the possibility that the distribution of stimuli across the different sleep stages would impact the results, we tested for differences in the relative number of stimuli in N3 (% stimuli in N3 relative to all NREM). We found no significant difference between conditions (long ISI: *median = 90.04 %, IQR = 7.16 %,* short ISI: *median = 91.06 %, IQR = 15.10 %*, long ISI>shortISI: *g = 0.20*, *p = 0.98, Wilcoxon signed-rank test*).

**Figure 2:**
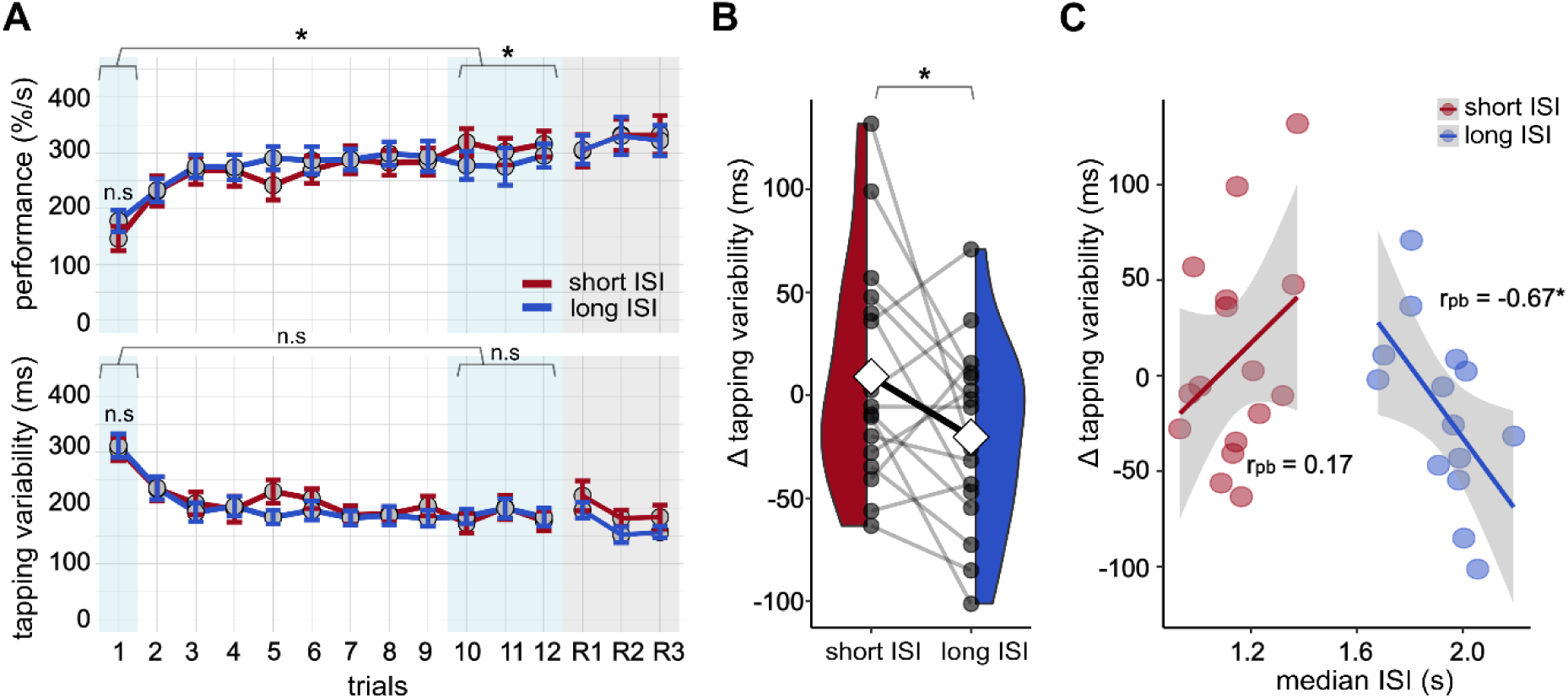
Learning and consolidation of the finger tapping task between long ISI and short ISI conditions. **(a)** Trajectories of performance and tapping variability across learning (1 — 12) and retrieval (R1 — R3) trials. Grey circles indicate means across participants and error bars indicate standard errors of the mean. Blue-shaded areas represent the baseline and learning plateau, and grey-shaded areas indicate retrieval. Asterisks represent significant differences with *p < 0.05; n.s. not significant; paired two-tailed Student’s t-test. **(b)** Overnight change in tapping variability was significantly different between short ISI and long ISI conditions. White diamonds indicate condition means, bold black line indicates mean difference, grey circles represent individual participants, and grey lines represent within-subject differences. **(c)** Overnight change in tapping variability in the FTT by condition and median ISI with linear model fit. The interaction of condition and median ISI was significant (β = −285.04, t = −2.013, p = 0.047). Robust percentage bend correlations indicated a significant association between ISI and overnight change in tapping variability only in the long ISI condition. Solid lines indicate the model prediction, grey shaded areas the 95 % confidence interval.

We fitted a robust linear mixed model for each behavioral outcome to assess whether the ISI manipulation within ON windows predicted the behavioral differences observed between the stimulation protocols. The model included median ISI, condition, and median ISI×condition as fixed factors and participant as a random intercept. Whereas the main effects of condition and median ISI alone did not significantly predict the overnight change in tapping variability (condition: *β = 398.95, t = 1.77, p = 0.08*; median ISI: *β = 155.11, t = 1.62, p = 0.13*), the interaction of condition and median ISI was significant (*β = −285.04, t = −2.01, p = 0.047*; **Figure 2C**). Robust percentage bend correlations within each condition suggested a significant association between ISI and overnight change in tapping variability only in the long ISI condition, where median ISI ranged from 1.68 s to 2.19 s (short ISI: *r_pb_ = 0.17, 95 % CI = (−0.35, 0.61), p = 0.53*; long ISI: *r_pb_ = −0.67, 95 % CI = (−0.89, −0.22), p = 0.02*). These findings suggest that longer ISIs predict the beneficial effects of the long ISI condition on overnight decrease in FTT tapping variability. However, ISI is not a strong predictor of behavioral response in the short ISI condition.

### Electrophysiological stimulus response

To explore the mechanism through which longer interstimulus intervals mediate improvements in overnight procedural memory consolidation, we quantified the stimulation response and its relationship to behavioral outcomes. A typical marker of stimulus response is the difference in low slow-wave activity between ON and OFF windows (ΔlowSWA). Whereas both protocols increased ΔlowSWA significantly (short ISI: *ΔlowSWA mean = 87.22* ± 146.55 *µV*^2^*, g = 0.57, p = 0.01,* one-sided one-sample t-test; long ISI: *ΔlowSWA mean = 94.87* ± *95.10 µV*^2^*, g = 0.94, p < 0.001,* one-sided one-sample t-test), they did not differ in their SWA response (*g = 0.11*, *p = 0.65,* two-sided Student’s t-test). A similar pattern emerged when assessing the adjacent SWA range (1.25 – 4 Hz; short ISI: *ΔSWA median = 9.63 µV*^2^ *(IQR = 21.54 µV*^2^*), g = 0.66, p < 0.001,* one-sided Wilcoxon signed-rank test; long ISI: *ΔSWA median = 7.77 µV*^2^ *(IQR = 11.73 µV*^2^*), g = 0.77, p < 0.001,* one-sided Wilcoxon signed-rank test; long ISI>short ISI: *g = −0.09*, *p = 0.62,* two-sided Student’s t-test). As the literature shows, ΔSWA is a coarse marker of response, which may not capture slow wave responses at different time scales^20,58^. Therefore, we decided to directly quantify K-complexes following auditory stimuli using a validated detection algorithm^42^.

The detection algorithm identified, on average, 9.8 (± 2.11) NREM sleep K-complexes per minute in the short ISI condition and 10.0 (± 1.86) NREM sleep K-complexes per minute in the long ISI condition, with no significant differences between the conditions. When restricting the analyses to different sleep stages or stimulated (first 2.5 hours of NREM sleep after first stimulus) and unstimulated parts of the night, we found a significantly increased K-complex density in N3 within the stimulated part of the night in the long ISI condition, but no other significant differences (**Table S1**). This finding might relate to a stimulation effect since most stimulations were applied during N3 sleep. On average, 299 (± 131.5) and 204 (± 90.8) K-complexes were associated with a stimulus in the short ISI and long ISI conditions, respectively. The average waveform of the detected stimulus-associated K-complexes showed a steep slow wave with a duration of ∼1000 ms and the characteristic P200, N550, and P900 components (**Figure S1**). The average time-frequency response relative to random NREM trials where no K-complex was detected was marked by a significant cluster of enhanced delta-activity coinciding with the wave’s up-to-down transition and a broadband decrease in high-frequency power. The findings confirm that, on average, the detected events align with the expected characteristics of a K-complex.

Stimulus-associated K-complexes were then compared between the long ISI and short ISI protocols. As displayed in **Figure 3A**, the relative number of stimuli followed by a K-complex was significantly larger in the long ISI condition than in the short ISI condition (*g = 0.70, p = 0.01*). The proportion of stimuli followed by K-complexes significantly predicted overnight reductions in tapping variability (*β = −3.72, t = −2.11, p = 0.045;* **Figure 3B**). From a stimulus-centered perspective, the proportion of stimuli followed by K-complexes reflects a tradeoff of the number of stimuli evoking a K-complex and the number of stimuli not evoking a K-complex. To further assess the individual contributions of these variables, we fitted a robust linear mixed model with main effects for the absolute number of stimuli with K-complexes and remaining stimuli without K-complexes. Both variables significantly contributed to the overnight behavioral change (**Figure 3C**): whereas the number of stimuli followed by a K-complex contributed to a reduction of overnight tapping variability (*β = −0.36, t = −2.22, p = 0.04*), the opposite was true for the number of stimuli not followed by a K-complex (*β = 0.11, t = 2.27, p = 0.03*).

**Figure 3:**
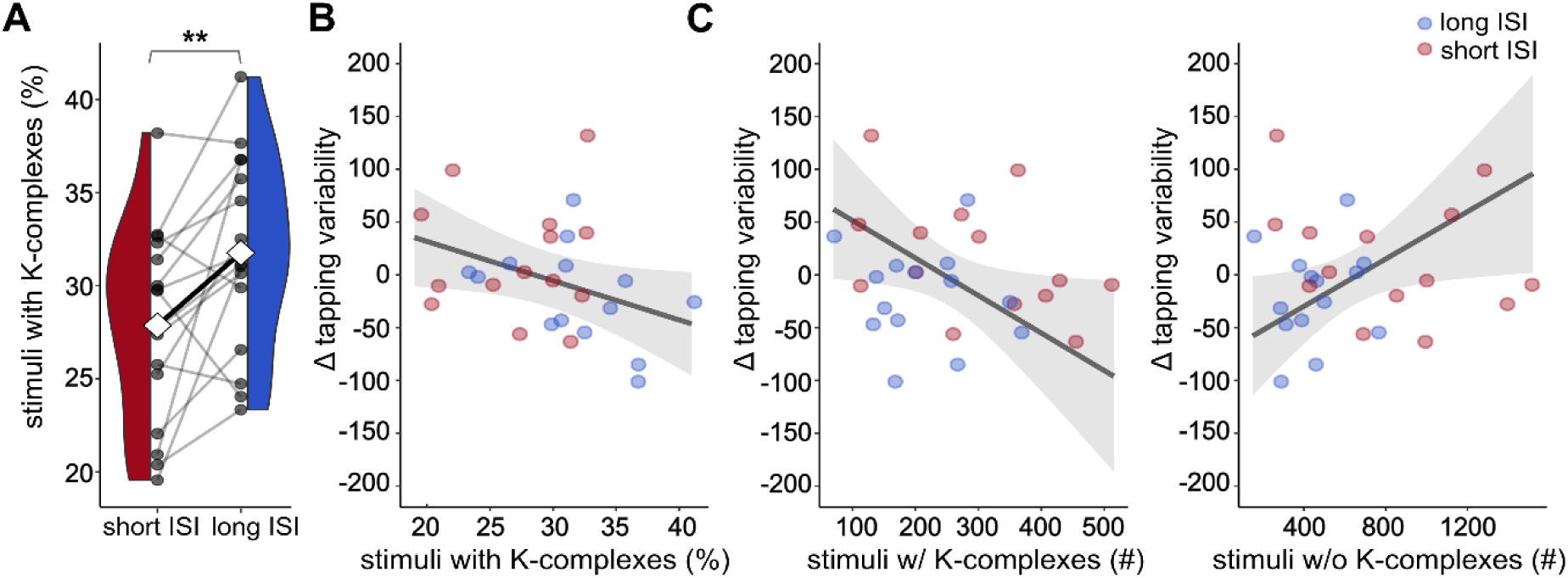
Proportion of auditory stimuli followed by detected K-complexes: comparison between short ISI and long ISI protocols and association with overnight change in tapping variability. **(a)** The proportion of stimuli followed by a K-complex was significantly higher in the long ISI than in the short ISI condition. Stimulus proportion per night was assessed by normalizing the total number of stimuli followed by a K-complex by the total number of ON-window stimuli administered during that night. White diamonds indicate condition means, bold black line indicates mean difference, grey circles represent individual participants, and grey lines represent within-subject differences. Asterisks represent significance levels with **p ≤ 0.01; paired two-tailed Student’s t-test. **(b)** Fixed effects plot of robust linear mixed model predicting overnight change in tapping variability in the FTT from the stimulus proportion followed by a K-complex. The effect of stimulus proportion followed by a K-complex was significant (β = −3.72, t = −2.11, p = 0.045). The solid line indicates the model prediction, grey shaded areas the 95 % confidence interval. **(c)** Fixed effects plot of robust linear mixed model disentangling contributions of stimuli followed by a K-complex and stimuli not followed by a K-complex to overnight change in tapping variability. The effects of stimuli with K-complexes and stimuli without K-complexes were significant (stimuli with K-complexes: β = −0.36, t = −2.22, p = 0.04; stimuli without K-complexes: β = 0.11, t = 2.27, p = 0.03). Solid lines indicate the model prediction, grey shaded areas the 95 % confidence interval.

Overall, the results suggest that long ISI stimuli evoke K-complexes more efficiently than short ISI stimuli. The balance between stimuli followed by a K-complex and remaining stimuli without a K-complex response may contribute to the behavioral effects on FTT tapping variability. Of note, the short ISI condition had both significantly more stimuli with K-complex responses and stimuli with no K-complex responses compared to the long ISI condition (**Table S1**), further substantiating that the absolute number of evoked K-complex responses is less crucial to behavior than their proportion to stimuli with no K-complex response.

### Relationship of interstimulus interval and electrophysiological stimulus response

We assessed the relationship between ISI and the stimulus-associated K-complexes to determine the optimal range of interstimulus intervals for evoking K-complex-like responses. To this end, a stimulus-associated K-complex was treated as a binary event, where each stimulus was either followed (1) or not followed (0) by a K-complex. The ISIs preceding the stimuli were assigned to approximately uniformly distributed ISI bins that were determined based on the distribution of ISIs across all nights (**Figure S2**). As displayed in **Figure 4**, the relative number of stimuli followed by a K-complex increased linearly across the lowest ISI bins and stabilized at ∼35 % starting at bin 4, spanning ISIs of 1.25 - 1.5 s. Individual lines showing the collapsed data from both intervention nights per participant highlight that all individuals followed a similar trajectory, with some variance in start and end levels. Overall, these results suggest that stimuli preceded by longer ISIs are more likely to induce K-complex-like responses than stimuli preceded by shorter ISIs. Moreover, the fact that the proportion of stimuli followed by K-complexes plateaus at ISIs > 1.25 s suggests an ISI threshold of > 1.25 s at which such a response should be evoked most reliably.

**Figure 4:**
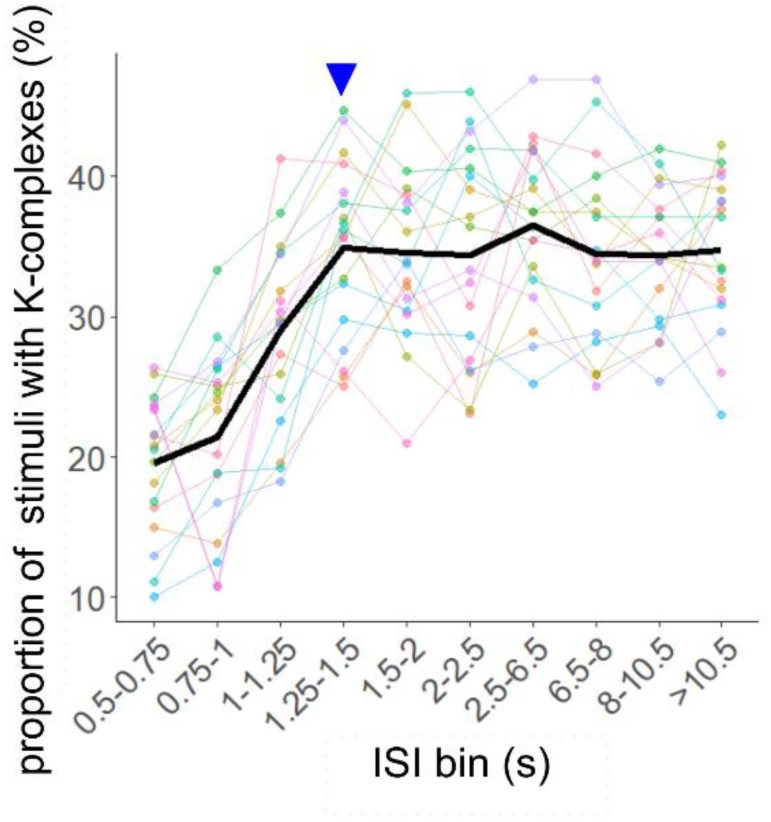
Proportion of stimuli followed by a K-complex by ISI bin. The normalization was performed with all stimuli per ISI bin, pooling short ISI and long ISI nights per participant. The solid black line represents the average across participants, colored lines represent individual trajectories. The blue arrow marks the bin at which the proportion of stimuli with a K-complex stabilized.

To test this prediction, we divided all stimuli into two categories: those preceded by an ISI > 1.25 s and those preceded by an ISI ≤ 1.25 s. Data were pooled across intervention nights for each participant. Averaging across stimuli from each category, we determined the AEP and time-frequency responses to stimuli in ON windows. Next, we contrasted the responses with average waveforms and time-frequency representations following comparable sham stimuli (i.e., meeting the same ISI criteria) in OFF windows (**Figure 5A**). Both stimulus categories evoked an AEP (**Figure 5B**), whose mean amplitude between 500 and 600 ms after stimulus onset showed a trend level difference between stimulus categories (*g = −0.26, p = 0.08*). The AUC of negative components between 200 and 1000 ms after stimulus onset was introduced to account for possible latency shifts in the N550 between participants. However, the AUC showed no significant difference between stimulus categories (*g = −0.06, p = 0.71*). The time-frequency response to supra-threshold (i.e., ISI > 1.25 s) stimuli revealed a positive cluster of activity immediately following the stimulus that extended from the low delta to the alpha frequency range and a significant power enhancement in a large sigma cluster starting at around 750 ms after the stimulus. In contrast, sub-threshold stimuli (i.e., ISI ≤ 1.25 s) showed no delayed sigma cluster but a similar early response stretching from the low delta to the alpha band. Sub-threshold stimuli also showed enhancements in the pre-stimulus delta band, which likely represents a response to the previous stimulus.

**Figure 5:**
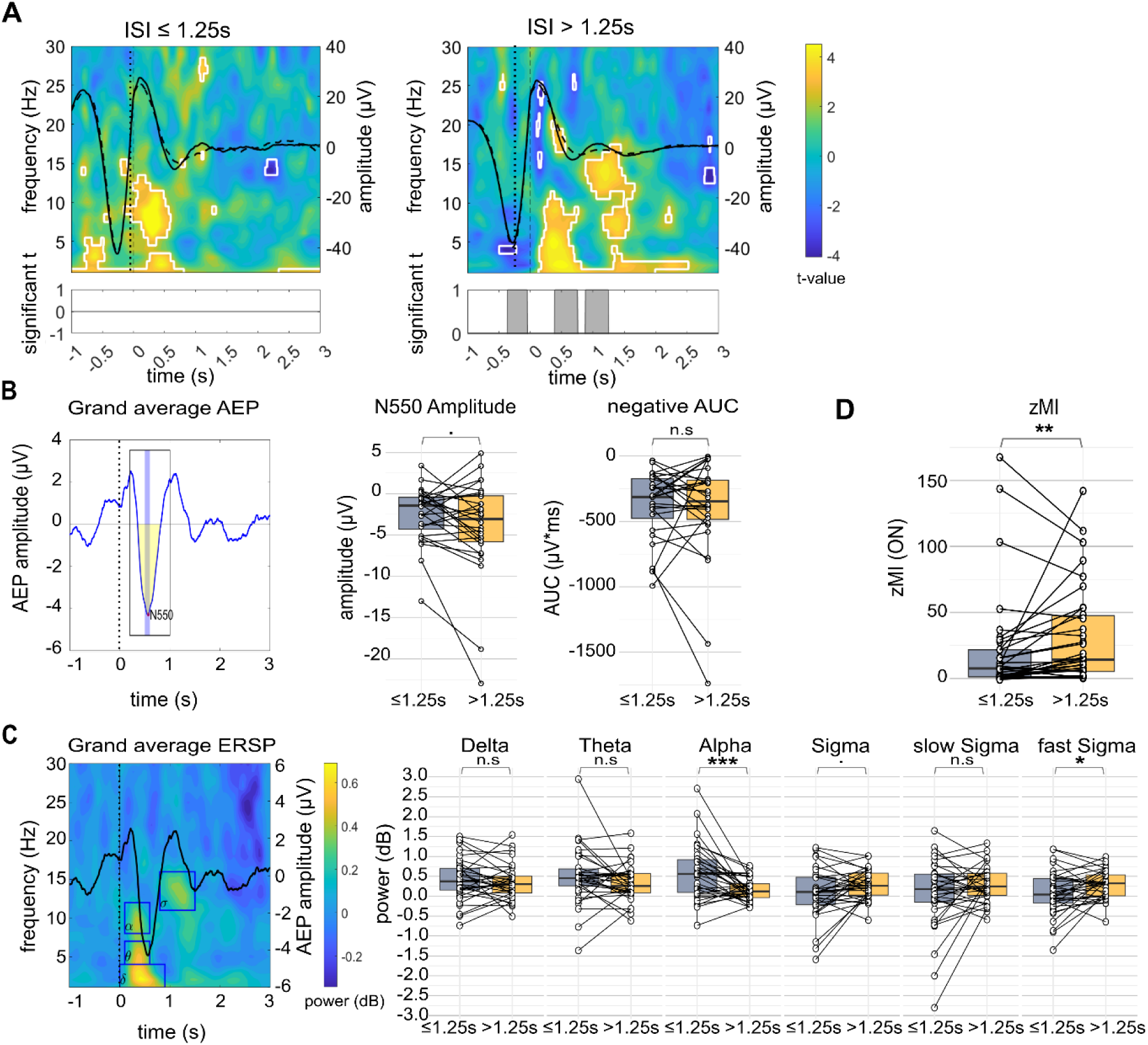
Responses to auditory stimuli categorized according to ISI cutoff. **(a)** T-maps relating to power in time-frequency space contrasting ON- and OFF-window stimuli classified by ISI. White margins indicate significant clusters after permutation-based cluster correction (paired two-tailed t-tests, α = 0.05). Solid black lines represent average waveforms of stimulus-locked responses in ON-windows. Dashed black lines represent average waveforms of stimulus-locked responses in OFF-windows. Significant differences between the two waveforms are marked in the bottom plot (paired two-tailed t-tests with permutation-based cluster correction, α = 0.05). Vertical dashed lines represent stimulus onset. **(b)** Left panel: Grand average auditory evoked potential (AEP) contrasting ON- and OFF-window stimuli with the N550 peak marked in red. The blue shaded area marks the time window considered for the average N550 amplitude. The black rectangle marks the time window considered for the negative area under the curve (AUC, indicated in yellow), reflecting the N550 AUC. Right panel: Comparison of amplitude and negative AUC extracted from regions of interest between stimulus categories. **(c)** Left panel: event-related spectral perturbation (ERSP) contrasting ON- and OFF-window stimuli, log-normalized by a whole-epoch baseline. The solid black line indicates the AEP (ON > OFF). Blue rectangles mark regions of interest. Right panel: Comparison of average power extracted from regions of interest between stimulus categories. The sigma band has been further subdivided into slow (< 13 Hz) and fast (≥ 13 Hz) component. **(d)** The z-scored modulation index (zM) between the phase of the 1 Hz oscillation and fast sigma power (≥ 13 Hz) in ON-windows compared between stimulus categories. **(b) – (d)** Boxplots show interquartile range and median. White circles refer to individual nights. Asterisks represent significance levels with *p < 0.05; **p ≤ 0.01; ***p ≤ 0.001, • trend (p < 0.1), n.s. not significant, paired two-tailed Student’s t-test or Wilcoxon signed-rank test.

To quantify the time-frequency responses of the two stimulus categories in more detail, we visually defined regions of interest based on the grand average ON-OFF time-frequency power representation and extracted average power values from these windows for each stimulus category and night (**Figure 5C**). This analysis confirmed what could be visually distinguished in the time-frequency plots: delayed sigma power showed a trend-level reduction in sub-threshold stimuli compared to supra-threshold stimuli (*g = 0.37, p = 0.05*). Splitting the sigma band into a fast (≥ 13 Hz) and slow (< 13 Hz) component, we found significant differences between stimulus categories in the fast (*g = 0.42, p = 0.03*) but not in the slow sigma band (*g = 0.32, p = 0.28, Wilcoxon signed-rank test*). Conversely, early alpha power was significantly more enhanced after sub-threshold stimuli compared to supra-threshold stimuli (*g = −0.78, p < 0.001, Wilcoxon signed-rank test*). We found no significant differences between stimulus categories in the early delta and theta bands (delta: *g = −0.13, p = 0.43*; theta: *g = −0.27, p = 0.21, Wilcoxon signed-rank test*).

Notably, the delayed sigma response was time-locked to the up-phase of the acoustically induced K-complex, implying the possibility that the phase-amplitude modulation between K-complex and sigma power was also part of the stimulus response. We therefore quantified the z-scored modulation index (zMI) between the 1 Hz slow oscillation, which falls into the frequency range of the K-complex, and fast sigma power (13 – 16 Hz) within 2 s after stimulus onset in ON-windows. The comparison of stimulus-associated modulation between the slow oscillation and fast sigma power showed a small but significant zMI enhancement after stimuli with a longer ISI compared to stimuli with a shorter ISI (*g = 0.24; p = 0.01, Wilcoxon signed-rank test*; **Figure 5D**).

## DISCUSSION

Here, we compared two automated up-PTAS protocols with systematic variations of the interstimulus interval regarding their effects on overnight procedural memory consolidation in the home setting. Compared to a protocol with shorter ISIs, the up-PTAS protocol with prolonged ISIs significantly reduced the tapping variability of a learned finger-tapping sequence after a single night of stimulation. These behavioral benefits were associated with the proportion of stimuli evoking a K-complex response, which is favored by longer ISIs. Specifically, we found that at ISIs > 1.25 s, a plateau is reached at which about 35 % of stimuli reliably evoke a K-complex. This response is also marked by an increase in fast sigma power, which is nested in the up-phase of the evoked K-complex. Although we performed this study in a less controlled setting, complete EEG data were recorded in 16 out of 18 participants (89 %), and complete EEG and FTT data in 14 out of 18 participants (78 %). These results suggest that

1. automated application of up-PTAS in naturalistic sleep in the home setting is both feasible and influences next-day behavioral responses,
2. up-PTAS can be optimized for overnight memory consolidation by introducing longer ISIs of at least 1.25 s,
3. long ISI up-PTAS favors K-complexes along with coupled fast spindles.

In support of our first hypothesis, the participants tolerated the mobile stimulation device well, with no drop-outs and good adherence to the sleep protocols. All but two participants completed tasks and questionnaires on time. In the short ISI condition, we achieved similar stimulation numbers as in a previous study using the same protocol in the laboratory setting^58^ and the electrophysiological responses we recorded replicated previous reports of PTAS-induced stimulation effects^5,20,59–61^. Similarly, our tablet-based FFT captured the stereotypical logarithmic learning curve observed in lab-based FTTs despite the less controlled setting^28,29,62–65^. Previous research has validated sleep stage and phase specificity of the *MHSL-SB*^66^ and replicated established electrophysiological stimulation effects with automated home-based PTAS in different study populations^2,5–7^. We extended these previous efforts by including an experimental comparison between PTAS protocols and a behavioral assessment of overnight procedural memory consolidation, which showed differential effects that were sensitive to the stimulation protocol.

Our second hypothesis predicted that the long ISI up-PTAS protocol would more efficiently drive memory effects than the short ISI protocol. Confirming this hypothesis, the long ISI protocol proved more effective despite involving significantly fewer stimuli. The overnight decrease in tapping variability observed in this study could be a readout of more fluent and stable motor performance after up-PTAS. This finding aligns with previous observations, according to which sleep selectively improved the speed of key press transitions that were the slowest before sleep^63^. Whereas we would have expected to find comparable effects of up-PTAS on performance, the observed pre-sleep differences in performance between the conditions limited our analysis. A detailed analysis dissociating accuracy and speed, which both contribute to the performance score, showed that pre-sleep differences were mostly accuracy-driven. Sheth et al. have argued that accuracy is influenced explicitly by task-related fatigue, causing it to reach an intermediate plateau level early during learning, whereas speed continues to increase over learning trials^67^. Therefore, it is conceivable that baseline fatigability in the FTT might have differed between the two assessment days, leading to performance differences, especially in the later learning trials.

In agreement with other studies rejecting the idea of a linear relationship between stimulus number and PTAS effectiveness^6,14^, the long ISI protocol produced better behavioral outcomes than the short ISI protocol despite applying significantly fewer stimuli. Our exploratory analysis suggests that the behavioral effect was mediated by the relative number of stimuli associated with a K-complex. The ISI determined the rate at which K-complexes occurred after stimuli, replicating earlier findings that showed more K-complexes after rare than after frequent stimuli^22,23,68^. The plateau proportion of stimuli evoking a K-complex at longer ISIs (> 1.25 s) was in a similar range (∼40 %) as reported previously, where single-frequency stimuli at slightly higher intensities and lower rates were employed^22,69^. Stimuli preceded by shorter ISIs (≤ 1.25 s) were associated with a lower probability of K-complexes and a trend towards lower N550 amplitudes on average. The stimuli that failed to evoke a K-complex response were negatively associated with overnight procedural memory retention relative to stimuli that did evoke a K-complex.

We could demonstrate that stimuli preceded by longer ISIs (> 1.25 s) produced a more pronounced spindle response than those preceded by shorter ISIs. Like K-complexes, spindles are known to have a refractory period, which has been estimated at 3 – 6 s^70^. Interestingly, stimuli presented at a longer latency of ∼2.5 s after a spindle, i.e., close to the end of the refractory period, significantly enhanced early spindle responses and memory retention compared to stimuli presented shortly after a spindle^70^. In our study, stimuli preceded by shorter ISIs might have fallen into the spindle refractory period, therefore failing to evoke a sigma response within the measured time window. Compared to stimuli preceded by longer ISIs, these stimuli also triggered an enhanced alpha response, which might relate to a higher number of stimulus-related microarousals. Together, our findings contribute to a body of research suggesting that incorporating longer ISIs in PTAS designs elevates the chances for stimulating optimal windows for memory consolidation.

Stimulus-evoked spindles, specifically fast spindles nested in the up-phase of the 1 Hz slow oscillation, seem likely to contribute to the sleep-dependent consolidation of explicit procedural memories. Indeed, previous studies have found functionally relevant enhancements in sigma activity^71,72^ and spindle characteristics, such as amplitude and density^73,74^, after motor sequence learning. Perturbation approaches have provided causal links between spindles and motor memory consolidation. For example, presenting odor cues that were selectively linked to the learned motor sequence in N2 sleep boosted memory retention while increasing spindle frequency and amplitude^75^. Furthermore, approaches selectively targeting sleep spindles with transcranial alternating current stimulation^76^ or auditory stimulation^18^ enhanced memory retention of learned motor sequences. In support of our findings, some reports highlight the role of fast spindles (≥ 13 Hz) in particular, in the context of motor sequence consolidation^57,74^.

Mechanistically, sleep is primarily thought to benefit consolidation of the effector-independent sequence information^77–80^, which is encoded in the hippocampus^81–83^. Fast spindles critically contribute to the neocortical-thalamic-hippocampal crosstalk^51^ that could be directly linked to memory replay. Coupling strength between spindles and slow oscillations, as assessed by the modulation index, has been linked with procedural memory consolidation^84^. Furthermore, studies employing simultaneous EEG and fMRI showed that spindles, particularly if they are coupled to slow oscillations, support the reactivation of task-relevant regions during sleep^64,65^. Spindle-related reactivation was positively correlated with overnight gains in task performance^64^. Of note, other studies using up-PTAS to enhance motor sequence consolidation have failed to observe behavioral benefits^16,85^. We speculate that insufficient fast spindle enhancement, imprecise slow-oscillation-spindle coupling, or stimulation during spindle refractory periods may have contributed to these negative findings.

### Limitations

While our results provide important insights into the mechanisms of PTAS and hopefully encourage more researchers to perform sleep studies in the home setting, we encountered some limitations that warrant careful consideration.

Firstly, whereas studies conducted in the home setting bear several advantages over classic laboratory studies, one difficulty is that not all behavioral tasks might be suitable in this setting. We especially identified task timing as an issue, as we had to exclude two participants from behavioral analyses for not adhering to the predefined schedule or mixing up tasks. Future studies should invest in designing well-standardized, automated tasks for different memory domains and implement better control and guidance mechanisms to ensure participants complete the correct tasks at the scheduled times.

Recording with a mobile device in the home setting allowed us to capture more naturalistic sleep but limited us to EEG data from only one frontal channel. Fast spindles tend to occur most frequently at central-parietal scalp locations^51,86,87^ but were also reported to be enhanced locally in those areas involved in task learning^65,71,73^. The fact that we still observed a fast sigma response with our limited setup could point to the prefrontal cortex being involved in task learning^88–90^. However, this interpretation would need to be supported by similar observations in other brain areas relevant to motor memory. The limited number of channels also hindered us from pursuing a spindle quantification approach due to small event numbers when detected with the *DETOKS* algorithm, which is, like most automatic spindle detectors, optimized for central channels^42,91,92^.

Secondly, we focused on comparing two active conditions, omitting a stimulation-free sham night to streamline the protocol. Thus, we were limited to comparisons between protocols or within-night ON-OFF windows. Although OFF windows might have been affected by activity from the previous ON window, potentially confounding ON-OFF contrasts, our findings align with the stimulation responses reported in other studies^5,20,59–61^. While we cannot entirely exclude possible negative impacts of stimulation on sleep architecture, participants’ sleep was within the normal range^93^, consistent with other studies that showed no effects of stimulation on sleep architecture^11,16^. Furthermore, it remains unclear if the short ISI condition failed to enhance motor memory consolidation or actively disrupted it, as both scenarios could have led to a significant difference between the short ISI and long ISI protocols. Supporting the first interpretation, previous PTAS studies have generally reported no effects on memory rather than detrimental effects^16,85^.

Thirdly, current K-complex detectors, including *DETOKS*, still offer limited precision and recall when considering human labels as the ground truth^42^. On the one hand, this suggests that some K-complexes might have been mislabeled (false positives) and, on the other hand, that some K-complexes were not detected at all (false negatives). Our data showed that the DETOKS tended to overestimate K-complex densities in N2 compared to previous reports^94^. However, as we do not expect any differential biases in K-complex detection between different ISI bins, we expect the trajectory of stimulus-associated K-complexes across ISI bins to be representative. We refrained from introducing more stringent criteria on duration, amplitude, or slope to maximize the number of detected K-complexes for statistical power. We would argue that even human labels, which are driven by morphological features, might not be the optimal ground truth, as K-complexes may change their morphology with increasing sleep depth^53^. Consequently, morphologically atypical K-complexes might be captured by data-driven algorithms like *DETOKS* but not by the human eye. Future improvements in detection approaches may circumvent these issues and improve detection accuracy.

### Conclusion

Extending studies that have suggested that a higher number of stimuli does not necessarily enhance PTAS effects, we find that a protocol with long interstimulus intervals was more effective in improving memory consolidation of a finger-tapping sequence than a protocol with shorter interstimulus intervals. Electrophysiologically, PTAS stimuli applied at longer ISIs drove K-complexes and slow-oscillation-spindle-coupling more efficiently than PTAS stimuli with short ISIs. The ability to perturb the system in favor of these responses could have advantages that extend beyond overnight memory consolidation. For example, K-complexes have been reported to regulate cardiovascular responses^95–98^ and positively affect heart function^99,100^. In a related context, K-complexes could also drive waste clearance through the glymphatic system via neurovascular coupling^101^. Importantly, our study shows that the long ISI PTAS protocol can be implemented effectively in the home setting, providing a framework for testing its broader application in longitudinal frameworks. While we could only look at variations in the ISI in this study, our understanding of factors influencing responses to PTAS is far from complete. Future studies may focus on additional factors that are likely to influence responses to auditory stimulation during sleep, like stimulus regularity and habituation effects or current arousal state.

## DATA AVAILABILITY

All data supporting the findings of this study are securely stored at servers of the University Children’s Hospital Zurich. Access and availability will be provided upon a material transfer agreement and after approval by the local ethics committee of the Canton of Zurich. Analysis code can be shared upon request.

## AUTHOR CONTRIBUTIONS

Data collection: VK, NM, NHM

Study design: VK, AM, RH

Methodology/ technical support: RS, NHM, MLF, WK

Formal analysis: VK, NM

Supervision: AM, RH

Writing – original draft: VK

Writing – editing: VK, SF, AM, RH

Writing – reviewing: NM, NHM, RS, MLF, WK

## ACKNOWLEDGEMENTS

We want to thank all the participants for their time and data contributions, Sven Leach for inspiring discussions and advice on the analyses, Frieder Kahl for programming one of the apps in our task battery, Lydia Kämpf for her active support during the initial set-up of the study, and all the lab members from the Huber sleep lab for their constant support and critical questions. Grammarly has been consulted for language editing of the revised manuscript.

