## Supplemental Material for "Longer Interstimulus Intervals Enhance Efficacy of Automated Phase-Targeted Auditory Stimulation on Procedural Memory Consolidation"

***Supplemental Material for Manuscript entitled "Longer Interstimulus Intervals Enhance Efficacy of Automated Phase-Targeted Auditory Stimulation on Procedural Memory Consolidation"***

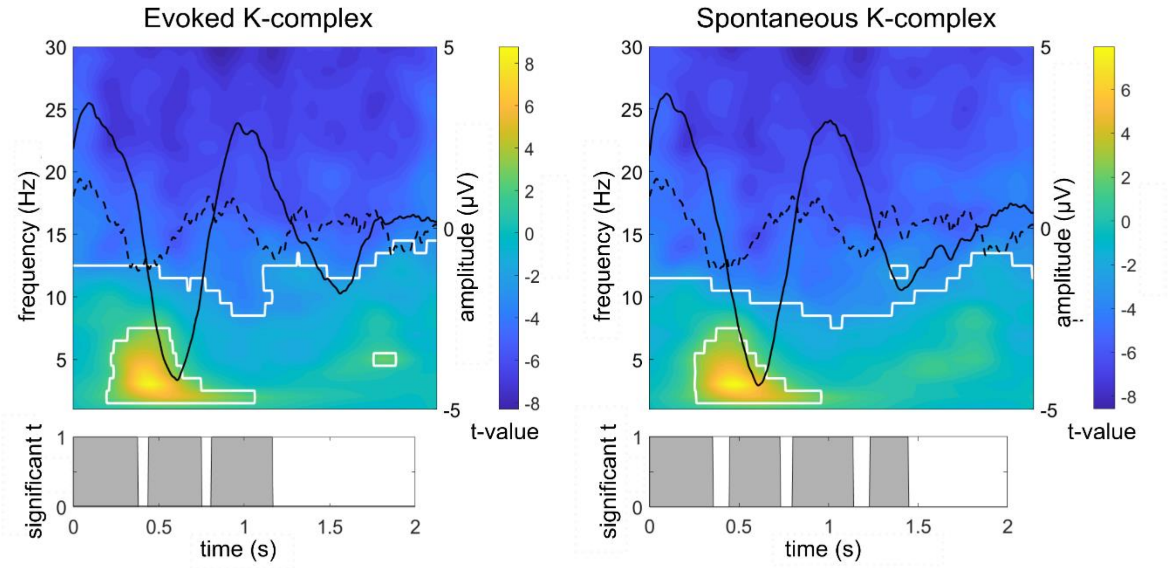


**Figure S1: Evoked (ON-window) and spontaneous (OFF-window) K-complexes detected by *DETOKS*.** Solid black lines represent average waveform across 2 s epochs locked to the starting point of detected K-complexes associated with ON-stimuli and sham stimuli in OFF windows. Dashed black lines represent average waveform across random 2 s NREM epochs not labeled as K-complexes. The bottom plot indicates areas where the difference between K-complex waveform and random NREM epoch waveform was significant after permutation-based cluster correction (paired two-tailed t-tests, α = 0.05). T-maps relate to power in the time-frequency space, contrasting K-complex epochs and random NREM epochs. White margins indicate significant clusters after permutation-based cluster correction (paired two-tailed t-tests, α = 0.05).

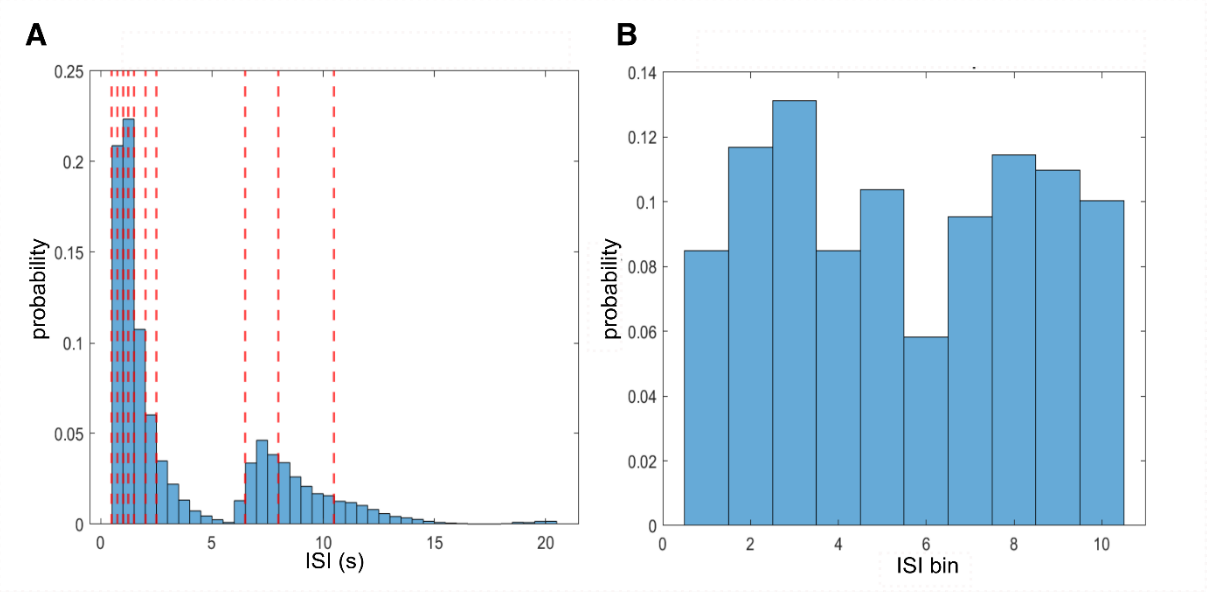


**Figure S2**: **Data-driven definition of ISI bins. (a)** Histogram of interstimulus intervals across the sample. Red dashed lines represent bin edges defined based on nine quantiles rounded down to the nearest 0.5 s, resulting in edges at 0.5 s (duplicate), 1 s (duplicate), 1.5 s, 2.5 s, 6.5 s, 8 s, and 10.5 s). The range between 0.5 and 2.5s was then subdivided into smaller steps for a more fine-grained parcellation, adding edges at 0.75 s, 1.25 s, and 2.0 s. **(b)** Histogram of ISI bin distribution across the sample.
