## Supplementary material for "Longer Interstimulus Intervals Enhance Efficacy of Automated Phase-Targeted Auditory Stimulation on Procedural Memory Consolidation": Table S1

**Table S1:** Comparison of detected K-complexes and stimuli associated with K-complexes between the stimulation conditions. Mean differences refer to the contrast longISI > shortISI. The reported p-values were obtained by paired Student's t-tests and effect size is reported in terms of Hedge's g. Significant comparisons are marked in bold. Abbreviations: N2 – NREM sleep stage 2, N3 – NREM sleep stage 3.

| | *shortISI (mean ± SD)* | *shortISI (min - max)* | *longISI (mean ± SD)* | *longISI (min - max)* | *Mean Difference* | *95% CI* | *p* | *g* |
|---|---|---|---|---|---|---|---|---|
| *NREM sleep K-complex density [1/min]* | 9.8 ± 2.11 | 5.9 – 12.8 | 10.0 ± 1.86 | 5.7 – 12.4 | 0.3 | [-0.57; 1.07] | 0.53 | 0.12 |
| *N2 sleep K-complex density [1/min]* | 7.0 ± 2.28 | 3.6 – 10.7 | 6.8 ± 1.95 | 3.2 – 10.0 | -0,1 | [-1.01; 0.75] | 0.75 | -0.06 |
| *N3 sleep K-complex density [1/min]* | 16.1 ± 1.81 | 12.6 – 18.5 | 16.6 ± 1.82 | 13.1 – 19.1 | 0.5 | [-0.40; 1.45] | 0.25 | 0.27 |
| *NREM sleep K-complex density 1st half [1/min]* | 11.8 ± 1.72 | 8.3 – 15.0 | 12.0 ± 1.97 | 8.6 – 14.8 | 0.2 | [-0.53; 0.89] | 0.59 | 0.09 |
| *N2 sleep K-complex density 1st half [1/min]* | 8.0 ± 1.71 | 4.8 – 11.9 | 7.3 ± 2.20 | 3.2 – 11.7 | -0.6 | [-1.50; 0.24] | 0.14 | -0.30 |
| *N3 sleep K-complex density 1st half [1/min]* | 15.6 ± 1.87 | 12.5 – 18.5 | 16.6 ± 1.98 | 13.1 – 19.4 | 1.0 | [0.23; 1.76] | **0.01** | **0.49** |
| *NREM sleep K-complex density 2nd half [1/min]* | 8.2 ± 3.17 | 3.1 – 12.6 | 8.3 ± 2.57 | 3.4 – 11.9 | 0.2 | [-1.22; 1.52] | 0.82 | 0.05 |
| *N2 sleep K-complex density 2nd half [1/min]* | 6.5 ± 2.48 | 3.0 – 11.3 | 6.7 ± 2.11 | 3.1 – 9.6 | 0.2 | [-0.82; 1.19] | 0.70 | 0.08 |
| *N3 sleep K-complex density 2nd half [1/min]* | 14.8 ± 6.35 | 0.0 – 21.3 | 15.6 ± 4.97 | 0.0 – 20.7 | 0.8 | [-1.62; 3.16] | 0.50 | 0.12 |
| *Stimuli with K-complex* | 299 ± 131.5 | 110 – 512 | 204 ± 90.8 | 46 – 369 | -94.3 | [-157.3; -31.4] | **< 0.01** | **-0.77** |
| *Stimuli without K-complex* | 800 ± 390.3 | 260 – 1516 | 434 ± 181.1 | 140 – 766 | -365.9 | [-530.2; -201.5] | **< 0.001** | **-0.96** |
| *Proportion K-complex evoking stimuli [%]* | 27.9 ± 5.27 | 19.6 – 38.2 | 31.8 ± 5.24 | 23.3 – 41.2 | 3.9 | [1.60; 6.74] | **0.01** | **0.71** |
